# Spectronaut-nf: A Nextflow Pipeline for Parallel Processing of DIA Data with Spectronaut

**DOI:** 10.64898/2026.07.29.741433

**Authors:** Chinmaya Narayana Kotimoole, Mohammad Arefian, Emma C. McKay, Sandeep Kasaragod, Grzegorz Skoraczyński, Ben C. Collins

## Abstract

**Summary:** Contemporary proteomics methods can now generate large-scale DIA datasets of thousands of files that demand substantial computational resources for efficient analysis. Spectronaut is a widely used platform for DIA data processing; however, large-scale searches are often constrained by computational performance and long execution times when run on single workstations. Here, we present Spectronaut-nf, a Nextflow-based pipeline that enables scalable and parallelized execution of Spectronaut analyses across high-performance computing (HPC) environments. The workflow divides directDIA analysis into modular stages, including spectral library generation, DIA searching, and merging results, allowing efficient distribution of tasks across multiple compute nodes. Benchmarking using 72 diaPASEF raw files using typical hardware demonstrated that Spectronaut-nf completed searches in 23.77 hours, compared with 39.09 hours on a Windows workstation and 67.04 hours on a single-node Linux HPC setup. Stress testing with 1,037 diaPASEF raw files further demonstrated the scalability and robustness of the workflow for large proteomics datasets. Across platforms, protein and peptide identifications remained consistent, with only minimal variability attributable to platform-specific differences. Overall, Spectronaut-nf provides a flexible, scalable, and efficient framework for high-throughput DIA proteomics analysis in HPC environments.

**Graphical Abstract:** 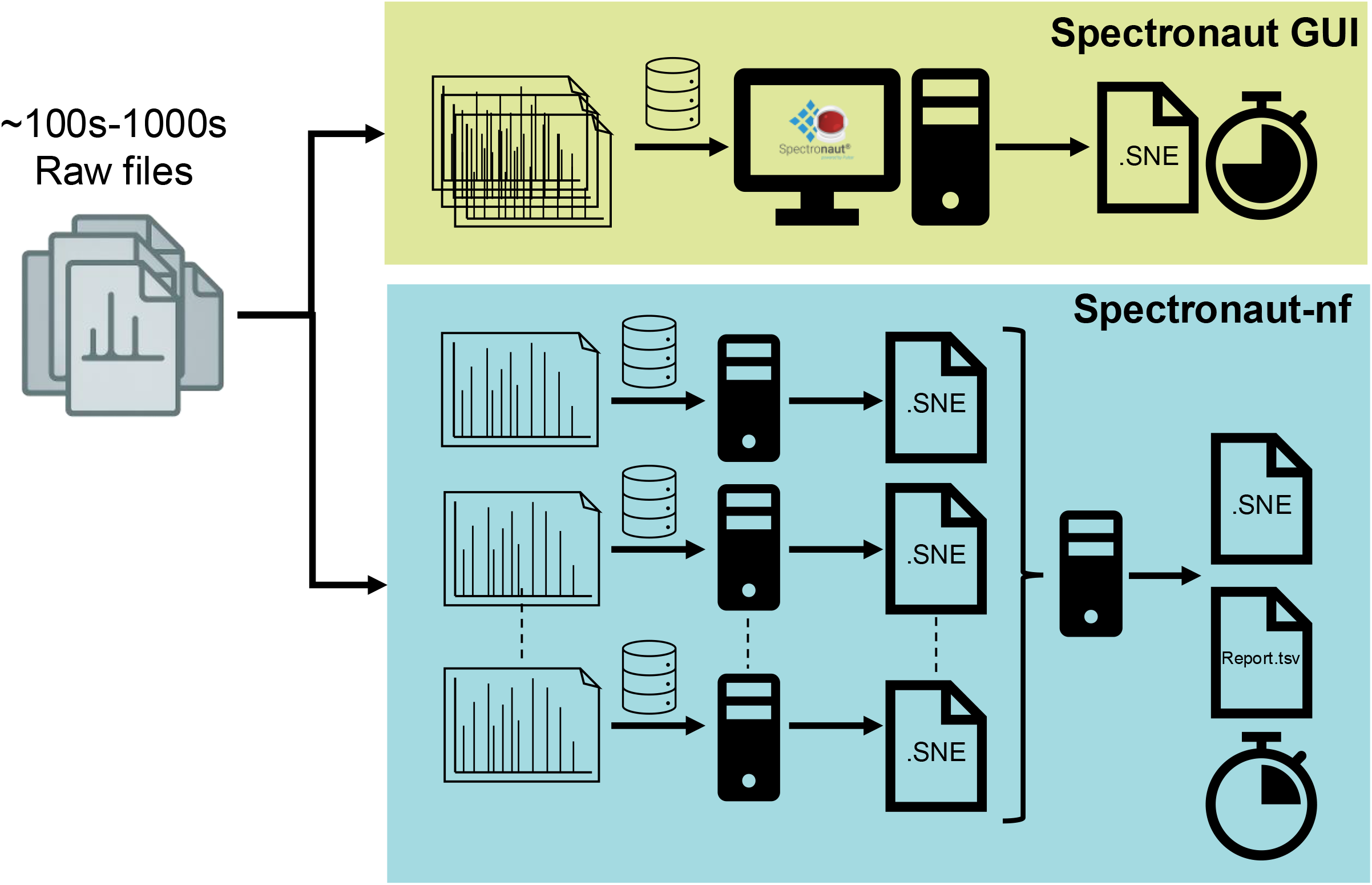

## Introduction

Data set size in contemporary proteomics is expanding rapidly with the introduction of automated sample preparation, fast and robust instrumentation that facilitates data acquisition at scale, and the need to generate very large sample sets that can address complex questions. In particular, projects focusing on population based clinical proteomics and perturbation/drug screens are pushing the boundaries with thousands to tens of thousands of samples analysed (1, 2). Strategies leveraging data independent acquisition (DIA) mass spectrometry now predominate in these large-scale studies. Analysis of DIA derived proteomic is computationally intensive typically requiring multiple CPUs and large memory space. Although newer versions of DIA analysis tools can process thousands to millions of spectra within minutes, availability/performance of computational resources and number of raw data files can be limiting factors (3). However, many researchers have access to institutional or national high-performance computing (HPC) resources and provision of an orchestration layer to facilitate analysis of DIA data at scale would be a useful addition to relieve such bottlenecks.

A broad variety of strategies and tools for the analysis of DIA data have been introduced (2,4–8). Spectronaut is a commercial tool prominently used by researchers in academic and industrial settings for analysis of DIA data in library-based or library-free modes (2,9). While Spectronaut has primarily been used in via graphical user interface (GUI) it includes a command line interface (CLI) and the recent introduction of Linux compatibility makes it more directly amenable to Linux-based HPC environments. An existing tool, quantms (10), that supports a broad variety of MS proteomics data analysis workflows can orchestrate parallel DIA data analysis on HPCs or cloud using DIA-NN (7), however, Spectronaut is not currently supported in this pipeline which primarily relies on open-source tools. We reasoned that a Nextflow (11) based pipeline would alleviate the bottleneck for analysing large-scale data sets via Spectronaut and be generally applicable to a variety of HPC or cloud based platforms.

## Experimental Section

### Spectronaut-nf implementation

Spectronaut-nf is implemented in Nextflow, designed mainly to enable parallel processing of large-scale datasets using Spectronaut in HPC and cloud platforms. The pipeline requires users to install or load Nextflow (https://docs.seqera.io/nextflow/install) and .NET v8.0 (https://dotnet.microsoft.com/en-us/download/dotnet/8.0) in their HPC/Cloud platforms. The users need to set the input parameters in *params*.*yaml* file, such as path to raw file directory and proteome database (FASTA), and optionally Spectronaut parameters for each stage where defaults are not used (.prop – can be generated in GUI). Where parallelization is used across compute nodes a floating Spectronaut license is required where the number of parallel instances of Spectronaut running across compute nodes should not exceed number of tokens on the license .

### Datasets

An in-house generated total proteome dataset from THP-1 cells was used for benchmarking and stress test of Spectronaut-nf workflow. Briefly, samples were prepared on an Opentrons OT-2 robot using Protein Aggregation Capture (PAC) with hydroxyl magnetic beads, enabling automated cleanup, digestion, and direct peptide loading onto Evotips (12). Peptides were separated using 40 samples per day (SPD) whisper zoom method of an Evosep One liquid chromatogram coupled to a timsTOF HT (Bruker Daltoniks, Bremen) in diaPASEF mode (12 x 2 variable window acquisition cycles) (13). In total, 205 diaPASEF raw files were used in the current study. Out of which, 195 were acquired with variable window diaPASEF, and 7 files using ion mobility-gas phase fractionation (IM-GPF) and 3 files from a longer gradient Whisper 20 SPD method were included (14). A detailed method description is included in Supporting Information. For comparison across platforms, we used 75 diaPASEF files. To evaluate the scalability of Spectronaut-nf, we constructed an enlarged benchmark dataset by duplicating files comprising 1,037 diaPASEF raw files and analyzed it using seven parallel compute nodes. The dataset consisted of 928 replicated experimental diaPASEF raw files generated from the original 96 raw files, together with 99 additional diaPASEF acquisitions from the same THP-1 samples and 10 gas-phase fractionation (GPF) raw files.

### Computer platforms

Windows platform: The directDIA search in Spectronaut GUI platform was carried out in a workstation with Windows 11. The workstation configuration was Intel Core i9-10900K (3.70GHz) with 20 CPUs and 128 GB of memory. However, only 18 CPUs were assigned in Spectronaut for the searches.

HPC platform: We used Kelvin2 research clusters hosted by NI-HPC centre (https://nihpc.github.io/nihpc-documentation/Kelvin2%20Hardware/). Spectronaut (v20.4) was used with Microsoft.NETCore.App (v8.0.5) in Rocky Linux (version 8.10) of Kelvin2 platform. All the searches were carried out using 96 (786GB/1TB memory) and 8 High Memory Dell PowerEdge R6525 nodes (2TB RAM). Both type of nodes consisted of 128 core AMD EPYC 7702 dual 64-Core Processors.

## Results

To enable the analysis of large-scale data sets via Spectronaut in HPC environments we developed a Nextflow pipeline (**Figure 1a**). The modular design facilitates the efficient allocation of resources on the HPC with library/DIA analysis steps running on raw data files being parallelized on different compute nodes whereas merging/combination steps running on a single node where high memory can be requested as needed (**Figure 1b**). Here, the direct DIA workflow of Spectronaut is split into 6 steps, *i) Library generation - Pulsar Stage 1*: Here raw files are searched in small batches in parallel against proteome database using the Pulsar search engine to generate intermediate search archives (.psar); *ii) Generate QSP*: All the .psar from previous step are integrated to generate a .qsp file harmonising the scoring across intermediate search archives; *iii) Library generation - Pulsar Stage 3*: Search Archives (.psar) from step i and corresponding raw files along with .qsp file generated in step ii are subjected to Pulsar Stage 3 search to generate final search archives file (.psar); *iv) Combine PSAR*: All search archive files (.psar) from Pulsar Stage 3 search are combined into a single experiment-wide search archive (.psar) and a spectral library (.kit format); *v) DIA search*: experiment-wide spectral library (.kit) is used to perform DIA analysis of each raw file in small batches in parallel to generate .sne files; *vi) Combine/merge SNE*: all .sne files are merged to form an experiment level result file (.sne). This .sne file can this be used in Spectronaut GUI for visualization or report generation. For large datasets (>150 .sne files), the pipeline automatically switches to Combine SNE process to generate a Report.tsv file directly instead of a merged .sne file as larger .sne files become more difficult to manage in the Spectronaut GUI and require substantially higher memory (RAM) for loading and processing. However, users can set the maximum number .sne files to be considered for SNE merging. Each step was executed by different Nextflow module *i.e*., *SN pulsarStages, combine psar, SN dia* and *Combine/merge sne*. Spectronaut-nf includes additional features such as random sampling of raw files for library generation, raw file batch size selection in single process for parallelization across nodes, and exclusion of raw files based on name patterns for DIA search.

**Figure 1.**
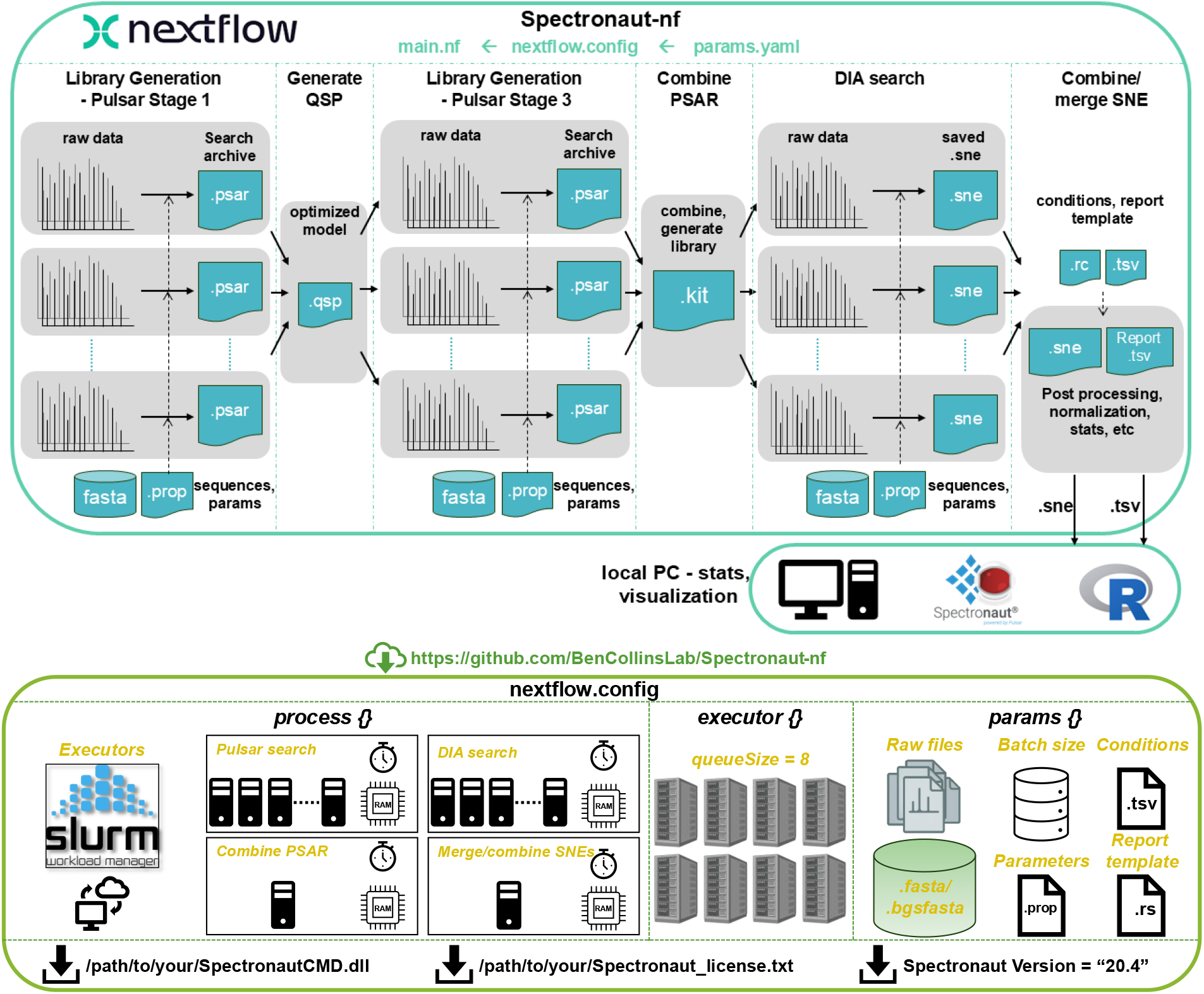
A schematic representation of Spectronaut-nf architecture.

To benchmark the workflow across platforms we tested Spectronaut-nf against typical ways of running the analysis using either the GUI on a Windows workstation or Linux CLI on a single node of our HPC using hardware typical for each workflow (**Figure 2a**). The direct DIA search of 72 diaPASEF files (288 GB) using Spectronaut in three modes i.e., Window GUI, single-node Linux CLI and Spectronaut-nf multi-node Linux CLI workflows completed in 39.09 h, 67.04 h and 23.77 h, respectively (**Figure 2b and Table S1**). As expected, the Nextflow-based Spectronaut-nf workflow finished the search ∼3x and ∼1.6x faster than the single-node Linux CLI and Windows GUI searches, respectively. This increase in the speed with Spectronaut-nf can be attributed mainly to parallel processing feature of Spectronaut-nf (**Figure S1**), which facilitates use of more CPUs across multiple nodes for the parallelizable steps of the workflow (see detailed info and execution timeline in **Table S1** and **Figure S1**).

**Figure 2.**
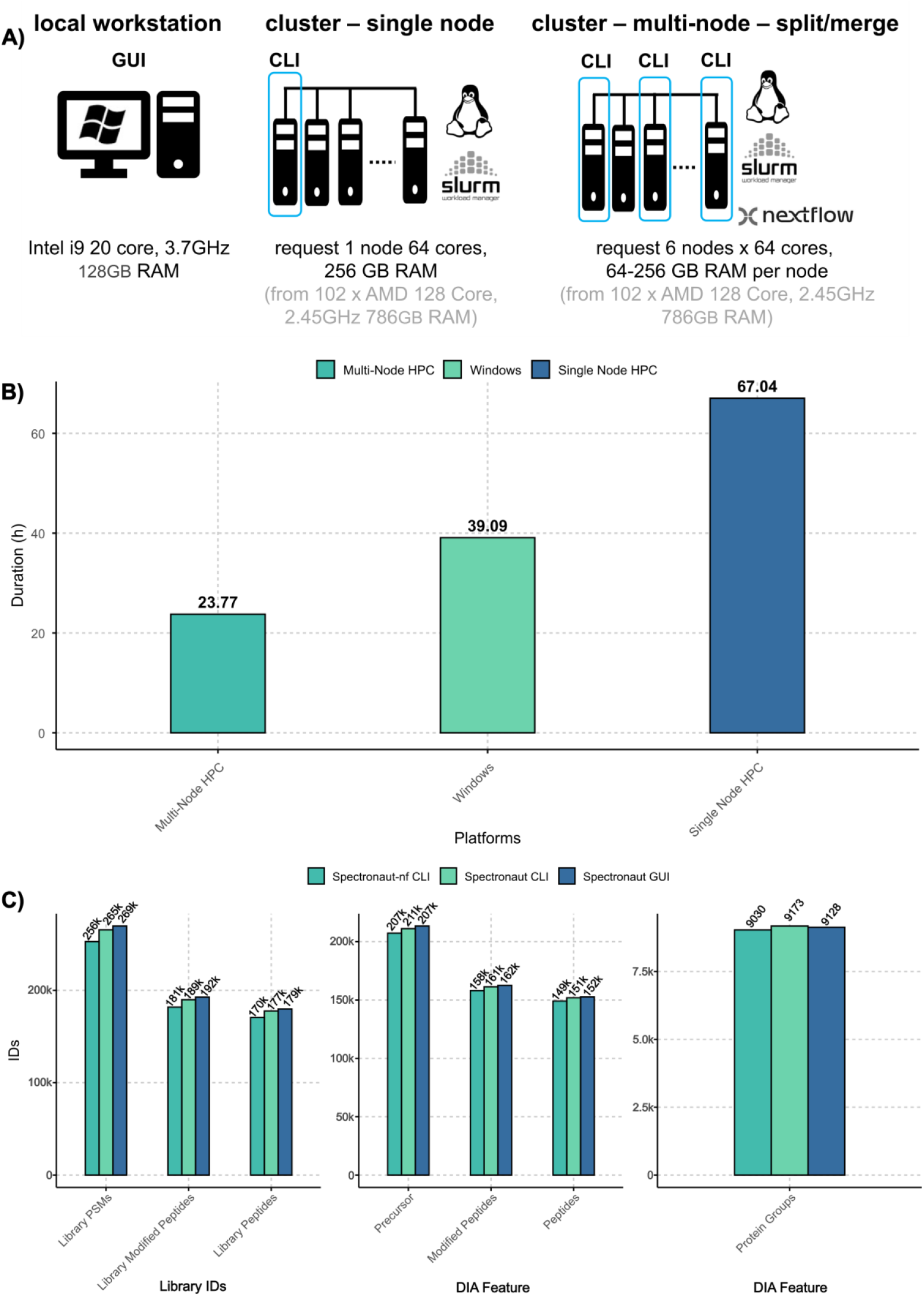
Total duration in hours taken by Spectronaut to search 72 diaPASEF raw files from 40 SPD runs across 3 computing platforms is represented as bar plot. c) Identifications at various levels for the 3 modes of running Spectronaut.

The library generation (Pulsar Stage 1, Generate QSP and Pulsar Stage 3) step for 72 raw files could be executed within ∼15.5 hours owing to *batch size* (i.e. number of raw files per batch) and *parallel processing* (i.e. number of parallel compute nodes) set to 2 and 6, respectively. Similarly, the *DIA search* step for 72 raw files was completed within ∼4.75 h owing to *batch size* and parallel processing of 4 and 8 within (**Figure S1**). The process-wise average of execution time, CPUs, memory and virtual memory from this benchmark experiment of Spectronaut-nf is available in **Table S1**.

These benchmarking searches resulted in the identification of more than 207,000 precursors, 158,000 peptides and >9,000 protein groups in all three platforms. However, across platforms we observed somewhat varying number of IDs at the precursor, peptide and protein group levels with Spectronaut-nf being modestly more conservative (**Figure 2c and Table S2**). In library generation we observed differences of 4 to 6% (PSMs, Modified Peptides and Peptides) and in DIA analysis 1 to 3% (Precursors, Modified Peptides, Peptides and Protein Groups) (**Figure 2c and Table S2**). These differences are caused among others by variations in floating-point arithmetic across different hardware platforms. Such variations introduce small numerical discrepancies in mathematical operations, which can accumulate over the course of an analysis and lead to noticeable result differences. However, despite these differences, the results remain functionally equivalent and remain under FDR control.

### Spectronaut-nf performance at scale

To examine the performance of Spectronaut-nf at scale we analyzed 1,037 diaPASEF files. The total execution time was 149.65 hours (i.e., 6 days, 5 hours, 41 minutes). A segmented bar plot shows the total duration taken to process all the input data in each task (**Figure 3**).

**Figure 3.**
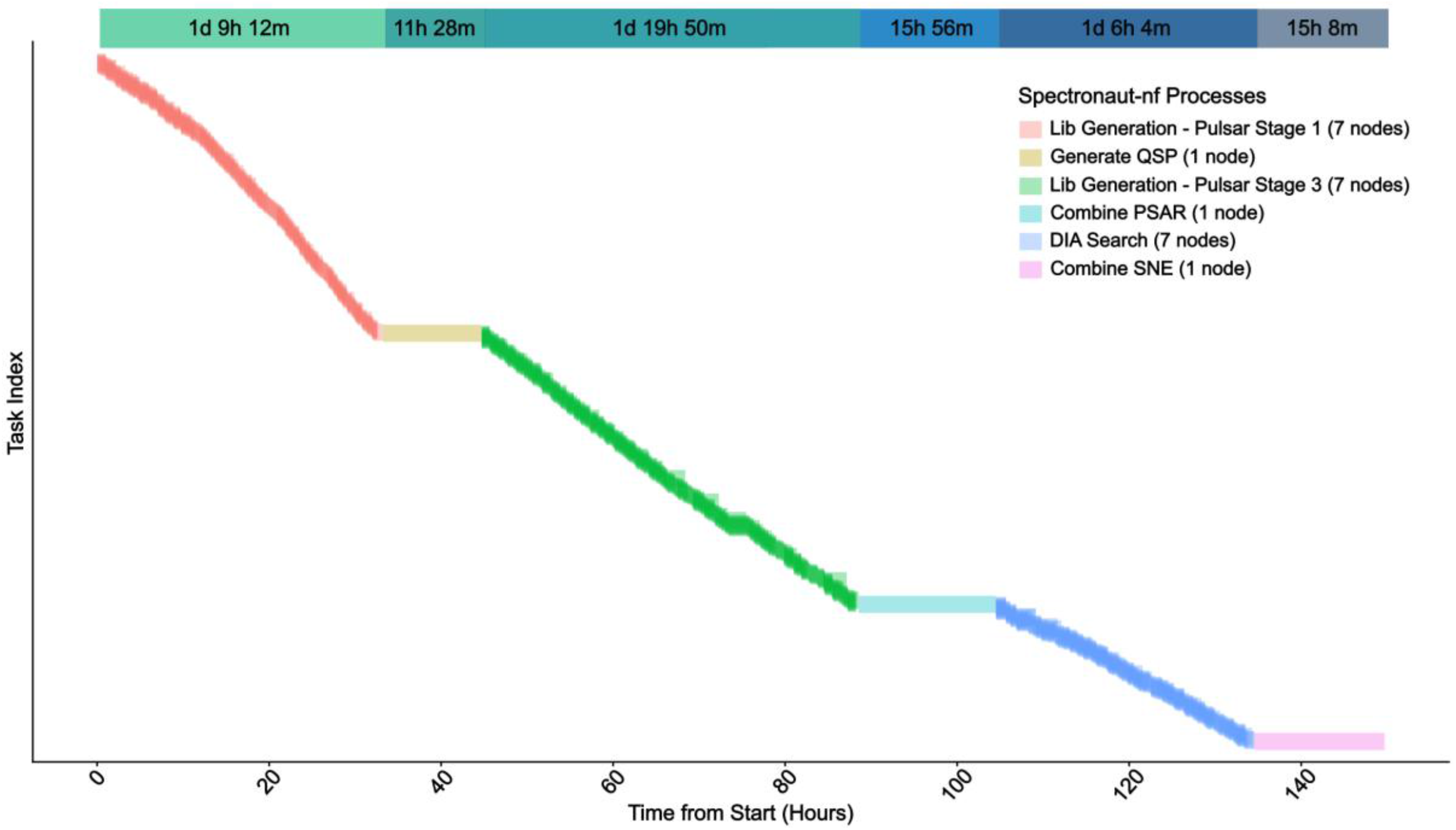
The execution time (queue waiting time + real time) of individual tasks across 6 stages (processes) of Spectronaut-nf pipeline for 1,037 dia-PASEF raw files were plotted to show the total execution time. Multi-node segments (1, 3, and 5) are overlaid with alpha factor. The segmented bars on top of the plot represents the time taken to execute each process for all the raw files.

On an average, library generation from a batch of 4 raw files took 0.86 and 1.15 hours of execution time in Pulsar Stage1 and 3, respectively, while DIA searches took 1.55 hours for a batch of 8 raw files (**Figure S2**). However, the HPC waiting time can vary based on batch size of raw files, amount of memory and number of CPUs are requested for each process. The single process memory-intensive tasks such as, generation of optimized model (.qsp), combining search archives (.psar) to generate single study specific library and combine SNE files to produce single report from all 1037 raw file took 11.47, 15.94 and 15.13 hours, respectively (**Figure S2**). Number of CPUs, memory (GB) and run time (hours) used per process are plotted in **Figure S3** and **Figure S4**. Single node processes such as *Generate QSP* and *Combine PSAR* consumed 122.7 GB and 129.4 GB (**Figure S3 and Figure S4)**.

## Conclusions

The Spectronaut-nf pipeline offers a robust, scalable, and flexible solution for high-throughput DIA data analysis that can scale to thousands of input files. By leveraging the Nextflow framework, it enables efficient parallel processing, integration with diverse computing environments, and streamlined management of complex tasks. The pipeline is freely available and open source on GitHub at https://github.com/BenCollinsLab/Spectronaut-nf.

## Data Availability

The mass spectrometry proteomics data have been deposited to the ProteomeXchange Consortium via the PRIDE (15) partner repository with the dataset identifier PXD078858.

## Supporting Information

Table S1: Process-wise resource usage by Spectronaut-nf from benchmark search

Table S2: Features identified from benchmarking DIA searches using Spectronaut across three computing platforms

Figure S1: Spectronaut-nf execution timeline for 72 dia-PASEF raw files

Figure S2: An average of total duration and real execution time from all the task in each process of Stress Test

Figure S3: A box plot representing CPUs, memory usage and execution time (Realtime) for all tasks from Stress Test across processes

Figure S4: A bar plot representing CPUs, memory usage and execution time (Realtime) for all tasks from Stress Test across processes

## Acknowledgements

We acknowledge institutional funding from Queen’s University Belfast to B.C.C. and BBSRC/MRC Prosperity Partnership (BB/Y00325X/1). We thank Tejas Gandhi, Oliver Bernhardt, and Monika Puchalska (Biognosys) for helpful discussions. We also gratefully acknowledge the members of the NI-HPC team, including James McGroarty, David Smith, and Vaughan Purnell, for their continuous support in assisting with implementation on the Kelvin2 HPC.

## Author contributions

C.N.K. and B.C. conceptualized and designed the study; C.N.K. and S.K. developed the pipeline; C.N.K. and M.A. and E.M. carried out the searches, preliminary data generation and writing-original draft; G.S. assisted with running Spectronaut for Linux and troubleshooting. All authors have read, contributed the editing of the manuscript and approved its contents.

