## Supporting Information for "Spectronaut-nf: A Nextflow Pipeline for Parallel Processing of DIA Data with Spectronaut"

**TABLE OF CONTENTS:**

**SUPPLEMENTARY METHOD**

THP-1 Cell Culture, MZ1 Treatment and Sample Preparation

**SUPPLEMENTARY TABLES**

Table S1: Process-wise resource usage by Spectronaut-nf from benchmark search

Table S2: Features identified from benchmarking DIA searches using Spectronaut across three computing platforms

**SUPPLEMENTARY FIGURES**

Figure S1: Spectronaut-nf execution timeline for 72 dia-PASEF raw files

Figure S2: An average of total duration and real execution time from all the task in each process of Stress Test

Figure S3: A box plot representing CPUs, memory usage and execution time (Realtime) for all tasks from Stress Test across processes

Figure S4: A bar plot representing CPUs, memory usage and execution time (Realtime) for all tasks from Stress Test across processes

**THP-1 Cell Culture and Sample Preparation**

Human monocytic THP-1 cells (ECACC #88081201) were cultured in RPMI-1640 (supplemented with 10% fetal bovine serum, 2 mM GlutaMAX™, penicillin–streptomycin, and ciprofloxacin) at 37 °C and 5% $CO_2$. Cells were lysed in sodium deoxycholate (1% SDC) lysis buffer containing TCEP and CAA, heated, sonicated, and clarified by centrifugation.

Proteomic sample preparation was performed using an Opentrons OT-2 liquid-handling robot following an automated Protein Aggregation Capture (PAC) workflow. Briefly, proteins were captured on magnetic hydroxyl beads, washed with acetonitrile, and digested using trypsin and Lys-C. Peptides corresponding to approximately 20,000 input cells were loaded onto Evotips and analyzed by LC-MS/MS using an Evosep One system coupled to a timsTOF HT mass spectrometer operated in dia-PASEF mode.

**LC-MS/MS analysis**

In brief, peptides loaded on to a Evotip were separated on an Evosep One system using the 40 SPD whisper zoom separation method. Peptides eluted from the Evotip were loaded on to an in-house analytical column (15 cm × 75 µm ID integrated emitter column from MSWill was packed with 1.5 µm Reprosil Saphir beads from Dr. Maisch). Using the integrated gradient method, peptides were separated based on their hydrophobicity by passing Mobile Phase A (0.1 % formic acid in Milli-Q water) and Mobile Phase B (0.1 % formic acid in acetonitrile). The analytical column was held at 50°C in a Bruker Column Toaster prior to the ionization.

Data were acquired on a timsTOF HT (Bruker Daltonics, Bremen, Germany) in dia-PASEF mode, employing 12 x 2 acquisition cycles comprising 24 variable m/z windows spanning 350–1,400 Da covering the ion mobility range (1/K0) from 0.64 to 1.37 V s/cm2. Peptides were ionizide in positive mode using CaptiveSpray ion source operated at 1,600 V capillary voltage, 3.0 L/min dry gas flow rate with a temperature of 180°C. Accumulation and ramp times were set to 100 ms. Collision-induced dissociation energy was ramped linearly as a function of the IM ranging from 59 eV at $1/K_0 = 1.6$ V s $cm^{-2}$ to 20 eV at $1/K_0 = 0.6$ V s $cm^{-2}$.

**Table S1: Process-wise resource usage by Spectronaut-nf from benchmark search.** An average execution time and computer resources used by each process of Spectronaut DIA pipeline in Spectronaut-nf from 72 diaPASEF raw file search

| PROCESSES | Batch size | No. of Tasks | Avg. execution Time | % CPU | Avg. CPUs | Avg. memory (RAM) | Avg. Virtual Memory (SWAP) |
|---|---|---|---|---|---|---|---|
| Library Generation (Pulsar Stage 1) | 2 | 32 | 0h:51m:29s | 2875.41 | 29 | 36.77 GB | 2.08 TB |
| Generate QSP (Pulsar Stage 2) | NA | 1 | 1h:33m:5s | 1021 | 10 | 32.70 GB | 2.15 TB |
| Library Generation (Pulsar Stage 3) | 2 | 32 | 0h:51m:29s | 2772.30 | 28 | 35.96 GB | 2.1 TB |
| Combine PSAR | NA | 1 | 3h:29m:58s | 175.20 | 2 | 32.10 GB | 2.15 TB |
| DIA search | 4 | 16 | 0h:51m:38s | 1563.76 | 16 | 40.31 GB | 2.06 TB |
| Merge SNE | NA | 1 | 3h:29m:56s | 96.2 | 1 | 48.50 GB | 2.05 TB |

**Table S2: Features identified from benchmarking DIA searches using Spectronaut across three computing platforms.** Number of precursors, peptides and proteins identified from directDIA search of 72 dia-PASEF raw files across Spectronaut GUI (Windows), Spectronaut CLI and Spectronaut-nf CLI (Linux) based search platforms

| **Name** | **Spectronaut GUI** | **Spectronaut CLI** | **Spectronaut-nf CLI** |
|---|---|---|---|
| Platform | Windows Workstation | Single-node HPC (Linux) | Multi-node HPC (Linux) |
| Library PSMs | 269,627 | 265,509 | 252,772 |
| Library Modified Peptides | 192,689 | 189,894 | 181,856 |
| Library Peptides | 179,815 | 177,635 | 170,679 |
| Precursor | 213,392 | 211,146 | 207,310 |
| Modified Peptides | 162,560 | 161,333 | 158,037 |
| Peptides | 152,693 | 151,871 | 149,116 |
| Protein Groups | 9,128 | 9,173 | 9,030 |
| Duration (h) | 39.09 | 67.04 | 23.77 |
| Elapsed Time | 1d 15h 5m 24s | 2d 19h 2m 24 s | 23h 46m 12s |

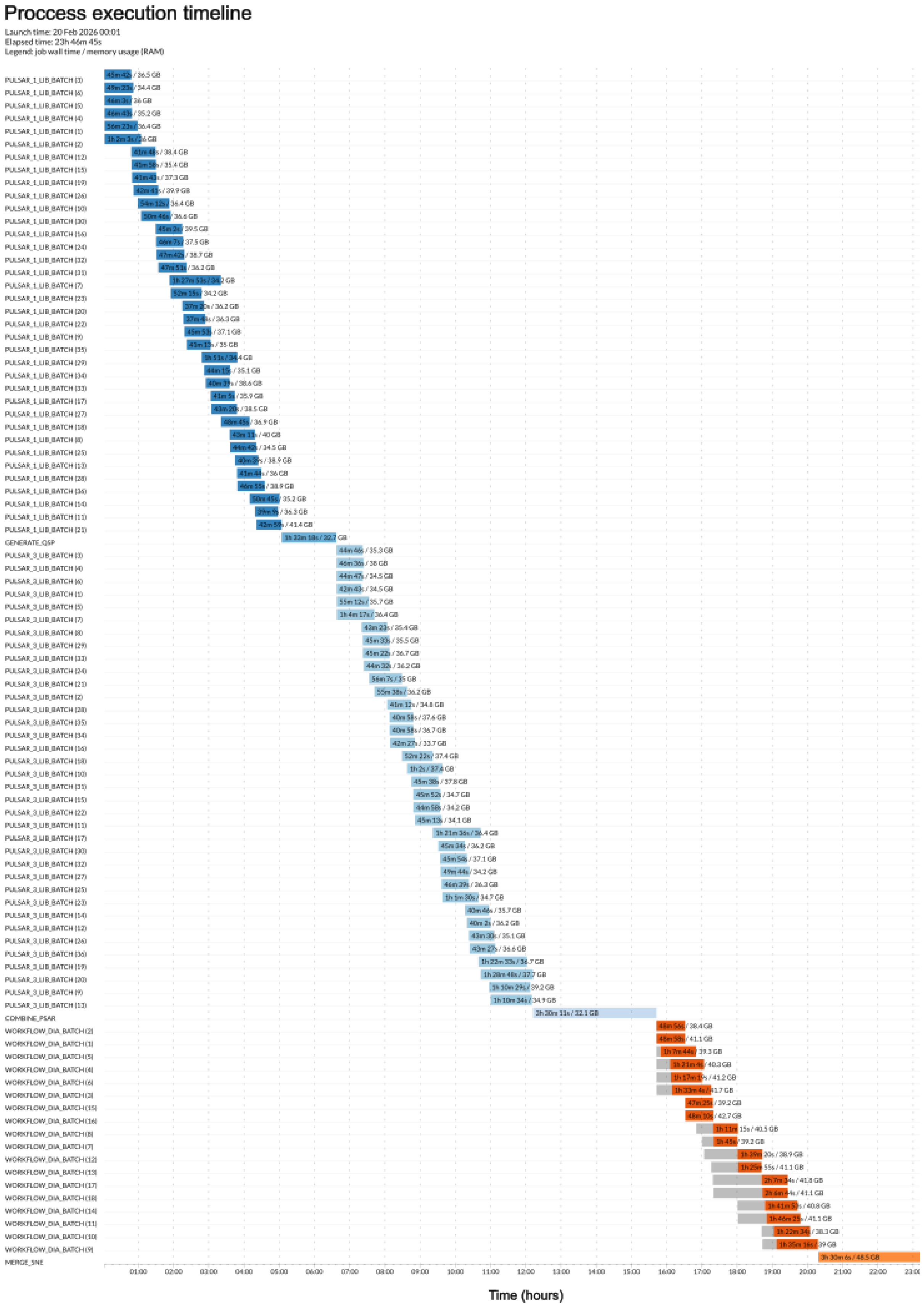


**Figure S1: Spectronaut-nf execution timeline for 72 dia-PASEF raw files.** A process execution timeline chart from the Spectronaut-nf analysis of 72 dia-PASEF raw file shows the total duration (queue waiting time + run time) and memory usage (RAM) of each process

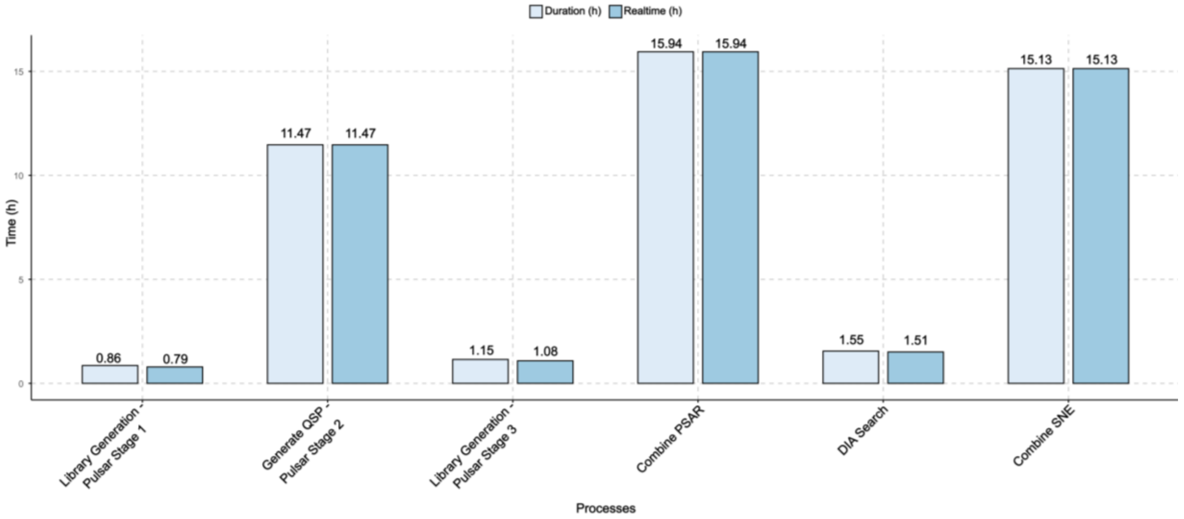


**Figure S2: An average of total duration and real execution time from all the task in each process of stress test.** The time taken by Spectronaut to complete a task (run time) with its waiting time in the queue (Duration=queue waiting time + run time) were stored in a trace file generated by NextFlow. These values for all the tasks of Spectronaut-nf Stress Test experiment were extracted, and processes-wise average was calculated and plotted.

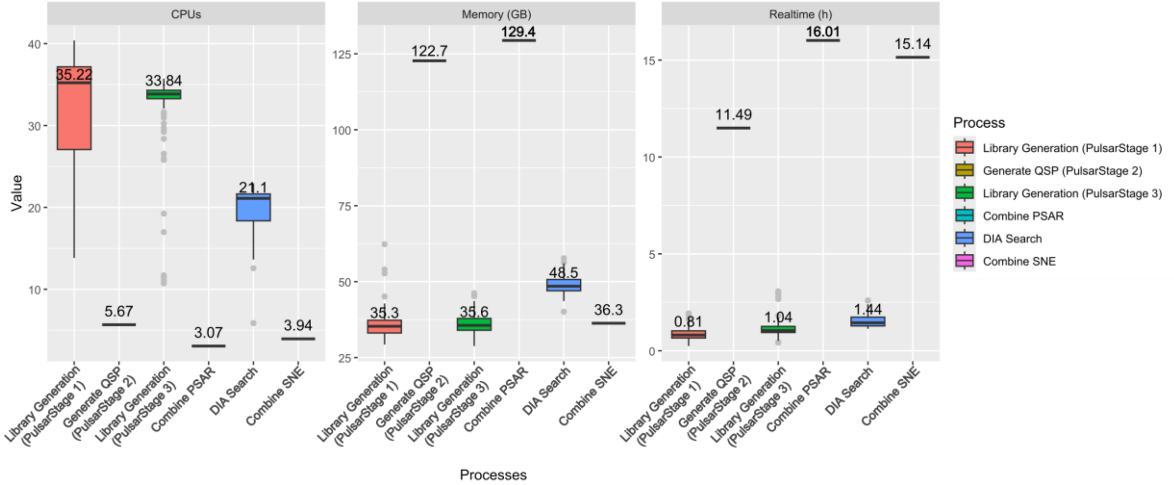


**Figure S3: A box plot representing CPUs, memory usage and execution time (Realtime) for all tasks from Stress Test across processes.** The CPUs, memory and execution time (run time) taken by all the tasks of Spectronaut-nf pipeline with 1037 raw file as part of Stress Test were extracted from trace file generated by NextFlow and plotted here process-wise. The median values from the CPUs, Memory used and real execution time taken in hours are highlighted next to the boxplot.

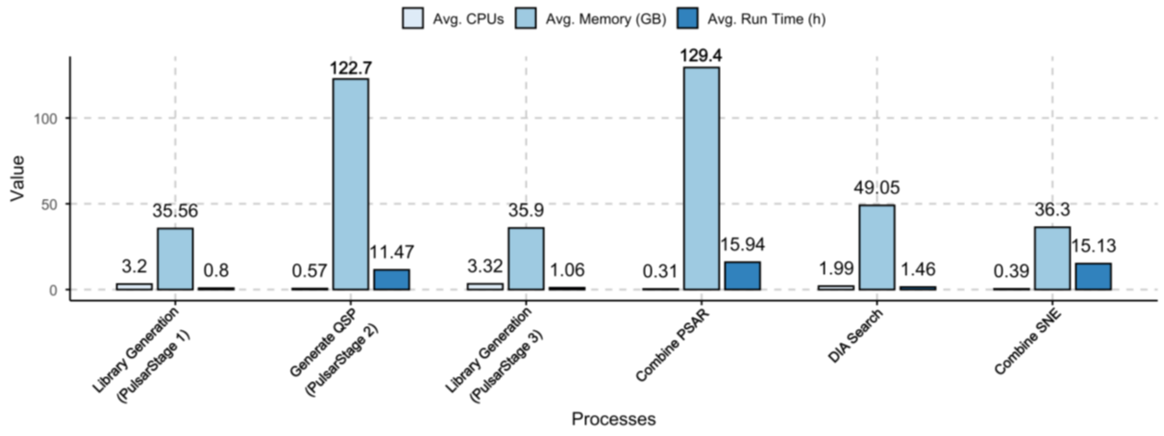


**Figure S4: A bar plot representing CPUs, memory usage and execution time (Realtime) for all tasks from Stress Test across processes.** An average of CPUs, Memory (GB) and real execution time (run time (h)) used by each process of Spectronaut-nf pipeline when 1037 raw files were searched are plotted. The average was calculated from the CPUs, Memory (GB) and run time (h) of each tasks stored in trace file of Nextflow were used.
